# Brain Functional Connectivity Signatures of Craving Across Substance Use Disorders: A Transdiagnostic Approach

**DOI:** 10.64898/2026.04.02.716016

**Authors:** Justin Böhmer, Luisa-Felicitas Esch, Katharina Eidenmueller, Richard O. Nkrumah, Lea Wetzel, Pablo Reinhardt, Norman Zacharias, Georg Winterer, Patrick Bach, Rainer Spanagel, Gabriele Ende, Wolfgang H. Sommer, Henrik Walter

## Abstract

Craving is a hallmark feature of substance use disorders (SUDs) and a major risk factor for relapse, yet reliable biomarkers that enable individual-level prediction remain scarce. Here, we applied connectome-based predictive modeling (CPM) to resting-state functional magnetic resonance imaging (fMRI) data in a transdiagnostic sample of individuals with cannabis, opioid, or tobacco use disorder (*n* = 78). Using CPM, we identified a distributed functional brain network that reliably predicted self-reported craving. Computational lesion analyses revealed key contributions from the right medial orbitofrontal cortex, right dorsal posterior cingulate cortex, and left lateral medial frontal gyrus. Importantly, the craving network generalized across two independent datasets. In alcohol-dependent patients (*n = 41*), the identified craving network, along with its positive and negative subnetworks, predicted distinct cognitive and motivational components of craving. In a second external dataset of smokers (*n* = 28), the craving network predicted both nicotine craving after abstinence as well as intra-individual changes in craving between sated and craving states. Together, these findings provide evidence for a robust, transdiagnostic craving signature in SUDs. Future work should assess the network’s predictive utility for longitudinal outcomes such as relapse risk and treatment response.

## INTRODUCTION

Craving, defined as an intense desire or urge to engage in specific motivated behaviors such as substance use, is a hallmark feature of substance use disorders (SUDs) and is strongly linked to poorer treatment outcomes and elevated relapse risk [1]. Consequently, identifying reliable biomarkers of craving has become a major goal in addiction neuroscience, offering the potential to guide personalized interventions and improve treatment success.

Functional neuroimaging studies have consistently linked craving to large-scale brain networks involved in salience detection, reward processing, attentional processing, memory functions, self-referential processing, and cognitive control [2–5]. However, most prior work has focused on group-level associations, limiting their translational potential for individual-level prediction. In recent years, there has been growing recognition that precision psychiatry requires methods that move beyond descriptive group comparisons toward models that generate accurate, individualized predictions of clinically relevant outcomes [6,7], as demonstrated in SUDs [8].

Connectome-based Predictive Modeling (CPM) is a data-driven machine learning approach that leverages whole-brain functional connectivity patterns to predict individual differences in behavior [9]. Unlike traditional univariate analyses, CPM capitalizes on distributed network information to identify predictive edges, construct summary connectivity scores, and generate robust predictions using cross-validation. CPM has been successfully applied to a variety of psychological and clinical outcomes, including substance use-related measures such as addiction severity in smokers [10], as well as markers of abstinence in opioid [11] and cocaine users [12].

In addition, four studies have applied CPM to investigate craving in SUDs [13–16]. Three of these studies examined craving within cue-reactivity paradigms, either using addiction-related stimuli [13] or mental imagery [14,16]. However, craving can also emerge spontaneously, independent of external triggers [4]. Resting-state functional magnetic resonance imaging (fMRI) offers a powerful tool to capture such intrinsic neural processes, providing insights into baseline craving propensity and trait-like vulnerability. One study employed CPM based on resting-state fMRI in individuals with opioid use disorder, identifying a brain network that predicts heroin craving as well as the change in heroin craving during abstinence [15].

Building upon this research, a critical next step is to establish predictive models in transdiagnostic samples that encompass multiple substances. Although different substances exert distinct pharmacological effects [17], evidence suggests that they share overlapping neural mechanisms [18–20]. A transdiagnostic modeling approach therefore provides a powerful means to identify common neural predictors of craving across SUDs, moving beyond substance-specific biomarkers toward shared circuit-level targets.

Consequently, in the present study, we applied CPM in a transdiagnostic SUD sample of individuals with cannabis, opioid, and tobacco use disorder to identify a functional connectivity network predictive of craving. To assess generalizability, we validated this network in an independent cohort of patients with alcohol dependence. Finally, we examined the sensitivity of the identified network to craving fluctuations in a second validation sample of tobacco users by testing whether it predicts intra-individual changes in craving from a sated to a craving state. By integrating transdiagnostic modeling, cross-cohort validation, and within-subject dynamics in craving, this study advances toward identifying a robust and generalizable connectome-based biomarker of craving in SUDs.

## MATERIALS AND METHODS

### Participants

A total of 91 non-detoxified individuals with SUD, aged 25–55 years, were recruited as part of a dual-center study conducted in Berlin and Mannheim. Thirteen participants were excluded due to incomplete craving data (*n* = 6) or insufficient fMRI quality (*n* = 7), resulting in a final sample of 78 participants (31 females). The mean age of the sample was 35.51 years (*SD* = 8.59). Sex was assessed by self-report and refers to assigned sex at birth. Participants were recruited from a predominantly ethnically homogeneous population; however, race and ethnicity were not formally assessed. Exclusion criteria included major psychiatric disorders (other than SUD), neurological disorders, and MRI contraindications. Inclusion criteria required participants to meet at least three diagnostic criteria for cannabis, tobacco, or opioid use disorder according to ICD-10. Diagnostic assessments covered both current (i.e., past 12 months) and lifetime substance use, including cannabis, tobacco, opioids, alcohol, amphetamines, ecstasy, cocaine, benzodiazepines, and pregabalin (Supplementary Table 1). The final sample comprised individuals with cannabis (CUD; n = 44), tobacco (TUD; n = 18), and opioid use disorder (OUD; n = 16). Participants were recruited through online advertisements. Subjects with OUD were recruited via outpatient substitution treatment clinics. All participants provided written informed consent prior to study participation. The study was conducted in accordance with the Declaration of Helsinki and approved by the medical ethics committees at both sites.

### Clinical measures

Craving was assessed using the Mannheim Craving Scale (MaCS), a reliable and valid self-report instrument designed to measure subjective craving during the past seven days across different substances [21]. The sum of the 12 MaCS items (rated 0-4) was used as the primary measure of craving. In addition, we assessed alcohol use using the Alcohol Use Disorder Identification Test (AUDIT), as well as smoking severity using the Fagerström Test for Nicotine Dependence (FTND). Depression, anxiety, and perceived stress were evaluated using the Beck Depression Inventory (BDI-II), the State-Trait Anxiety Inventory (STAI), and the Perceived Stress Scale (PSS), respectively.

### MRI data acquisition

All images were acquired on two 3T Siemens Magnetom Prisma MRI scanners in Berlin and Mannheim with a 64-channel head coil (Siemens Healthineers, Erlangen, Germany), respectively. For each participant, 500 functional images were acquired during resting-state using a gradient-echo echo-planar imaging (GE-EPI) sequence (TR = 869 ms, TE = 38 ms, 58° flip angle, 60 slices, 2.40 mm slice thickness, FOV = 210 mm, 2.4 x 2.4 mm^2^ in-plane resolution, bandwidth = 1832 Hz/Px, time of acquisition = 7:23 minutes). During data acquisition, participants were asked to lie as still as possible while keeping their eyes closed without thinking of anything specific or falling asleep. Additionally, high-resolution T1-weighted 3D structural images (MPRAGE, TR = 2000 ms, TE = 3.03 ms, 9° flip angle, isotropic voxel size of 1 mm^3^, bandwidth = 130 Hz/Px) and a B_0_-field map (TR = 698 ms, TE1 = 5.19 ms, TE2 = 7.65 ms, 54° flip angle, isotropic voxel size of 2.4 mm^3^, bandwidth = 279 Hz/Px) were acquired. <u>fMRI preprocessing</u> Resting-state fMRI data were pre-processed using HALFpipe [22], a containerized pipeline based on fMRIprep [23]. Pre-processing included motion correction, slice timing and distortion correction, ICA-based denoising (ICA-AROMA), spatial smoothing (FWHM = 6 mm), grand mean scaling (M = 10.000), temporal filtering (Gaussian-weighted high-pass width of 125 s), spatial normalization and registration to the MNI152 template. Manual quality control was performed to evaluate temporal signal to noise ratio, excessive movement, and quality of registration to the T1 image. Subjects with excessive head motion (mean FD > 0.5mm, or maximum FD > 4 mm, or > 30% of frames with FD > 0.5mm) were excluded from further analyses.

### Functional brain network construction

Functional brain networks were constructed by calculating the Pearson correlation between the time series of all pairs of regions defined by the Brainnetome Atlas [24], resulting in a 246×246 functional connectivity matrix for each subject. The resulting correlation coefficients were normalized using Fisher’s r-to-z transform.

### Connectome-Based Predictive Modeling (CPM)

We applied CPM [9] to identify functional connections associated with craving. In the training set, edge weights (i.e., functional connectivity values) were correlated with behavioral craving scores using Pearson’s correlation. Edges significantly associated with craving (*p* < .005) were selected to construct positive and negative predictive networks. The positive network included edges where stronger connectivity predicted higher craving, while the negative network comprised edges where weaker connectivity predicted higher craving. Each edge was assigned exclusively to one network, rendering the positive and negative networks mutually exclusive. These networks were then aggregated into a combined predictive network capturing both patterns of association.

Leave-one-out cross-validation (LOOCV) was used, with each subject left out once while the model was trained on the remaining data. For each subject, the strength of the positive, negative, and combined networks was calculated by summing the weights of the respective significant edges. These network strength values were entered into linear models to estimate craving. The resulting model coefficients were then applied to the left-out subject’s network strength values to generate predicted craving scores.

Model performance was evaluated by calculating the correlation between true and predicted craving scores using Spearman’s *rho* and Pearson’s *r.* Additionally, we computed the coefficient of determination (*q²*) and mean squared error (*MSE*). To assess statistical significance, craving scores were randomly shuffled 10,000 times, and CPM was rerun on the permuted data. For each permutation, Spearman’s *rho* between the true and predicted scores was computed. Spearman’s *rho* was used as it does not assume normality, making it more appropriate for skewed or ordinal data. *P*-values were then derived as the number of permuted *rho*-values greater than or equal to the true *rho*-value, divided by the number of permutations. One-tailed *p*-values were calculated, as only positive correlations between predicted and actual values reflect successful prediction.

### Control analyses

To evaluate the robustness of our findings, we tested both a more liberal (*p* < .01) and a more conservative (*p* < .001) significance threshold for feature selection. In addition, to account for potential confounders, we conducted sensitivity analyses by using partial correlations instead of full correlation during feature selection, controlling individually for substance class, age, sex, site, education, smoking severity, head motion, as well as collectively for all covariates.

### Computational lesion analysis

To examine the importance of each brain region for craving prediction, we performed node-wise and network-wise computational lesion analyses. For node-wise lesioning, each brain region (node) was systematically removed from the full 246 × 246 connectivity matrix, and CPM was rerun using the resulting lesioned matrix (245 × 245). For network-wise lesioning, nodes were grouped into eight large-scale canonical brain networks, based on the seven-network parcellation by Yeo et al. [25], with an additional subcortical network comprising all subcortical nodes. Afterwards, a given subnetwork (e.g., the visual network, consisting of 41 nodes) was removed from the overall network and CPM was re-run based on the lesioned matrix (e.g., 205 x 205). The difference in prediction between the true and lesioned model, computed using Steiger’s Z-test [26], indicated the importance for each brain region and network.

### External validation

For validation analyses, we tested the predictive performance of the identified craving network in two external datasets. First, to assess the generalizability for other substances, we leveraged a sample of male patients with alcohol dependence (*n* = 41) recruited from inpatient care [27]. Subjects underwent resting-state fMRI scanning and reported alcohol craving based on the German version of the Obsessive-Compulsive Drinking Scale (OCDS-G), which aligns conceptually with the craving measure implemented in our discovery sample [21]. The OCDS-G assesses alcohol craving across two subscales: The *Thoughts* subscale, reflecting cognitive preoccupation with alcohol, and the *Impulse to Act* subscale, capturing the urge to drink. Detailed sample characteristics can be found in the Supplementary Information (Supplementary Table 2).

Second, to assess the sensitivity of the identified craving network to craving state fluctuations, we examined an independent sample of subjects with tobacco use disorder (*n* = 28) who underwent resting-state fMRI scanning twice: once in a sated state at baseline, and once in an experimentally induced craving state after a minimum of 10 hours of smoking abstinence after baseline (see Supplementary Table 3 for sample characteristics). Craving was measured at both timepoints using the German version of the Questionnaire on Smoking Urges (QSU-G). The QSU-G measures nicotine craving across two factors: *Factor 1*, indicating the desire and intention to smoke, and *Factor 2*, assessing the anticipated relief from negative affect.

Resting-state fMRI data from external samples were preprocessed, quality-checked, and normalized analogously to the discovery dataset. For validation, we used the identified craving networks (positive, negative, and combined) as masks to extract connectivity strength for each subject in the external samples. These values were entered into a linear model to predict craving scores. We assessed prediction accuracy using Spearman’s *rho* and Pearson’s *r*. To test significance, we ran 10,000 permutations by randomly shuffling true craving scores and recalculating correlations. *P*-values reflect the proportion of permuted *rho*-values equal to or greater than the observed *rho*.

## RESULTS

### Sample characteristics

No significant differences in self-reported craving were found between SUD groups (*F*(2,75) = 2.08, *p* = .132). Craving did not differ significantly by site (*t*(76) = -0.34, *p* = .737) or sex (*t*(76) = -0.34, *p* = .737) and was not significantly correlated with age (*r* = -0.21, *p* = .065) or mean head motion (*r* = - 0.09, *p* = .493). See Table 1 for an overview of demographic and clinical variables.

**Table 1.**
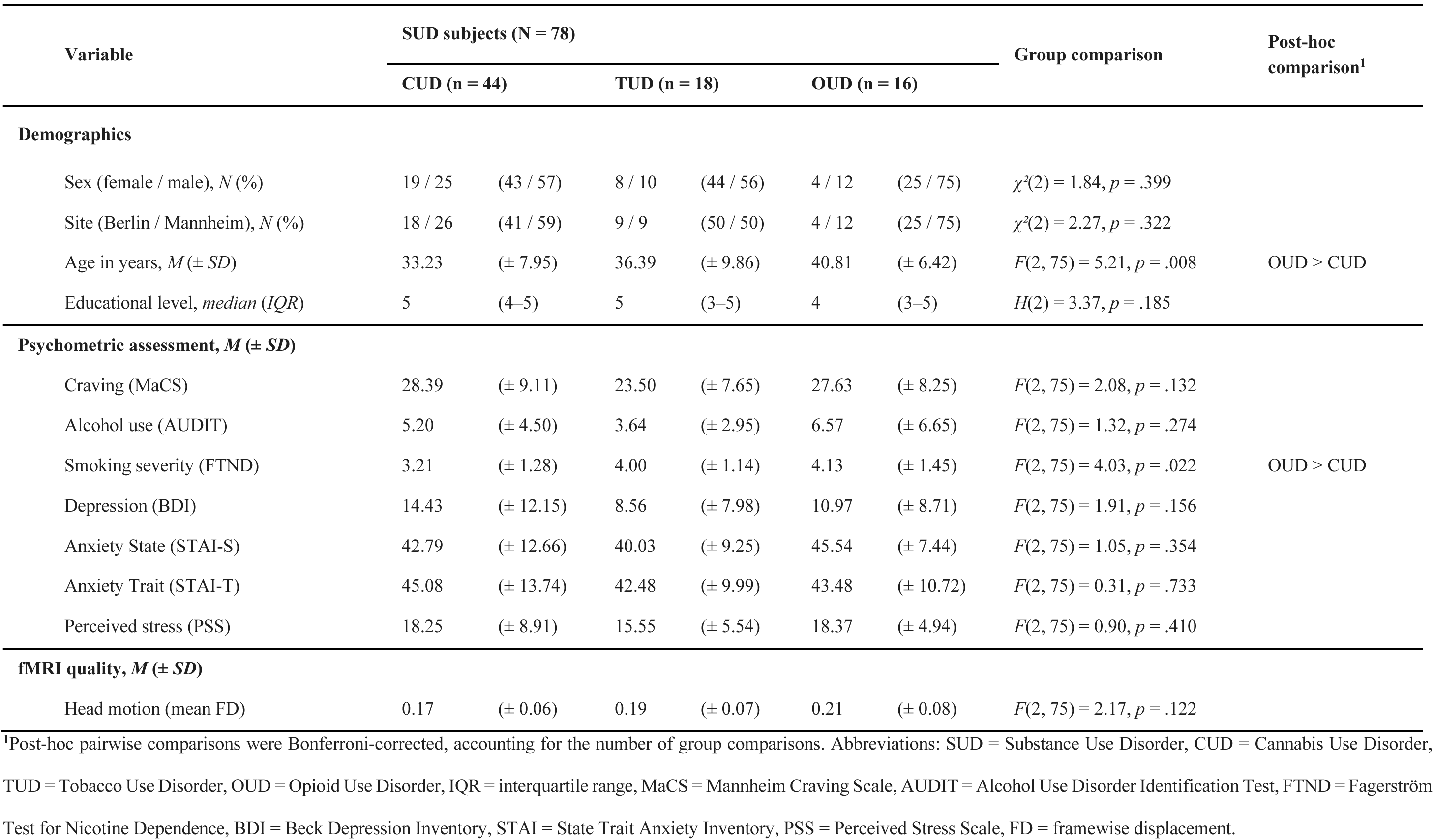
Sample description and demographics.

| Variable | SUD subjects (N = 78) |  |  |  |  |  | Group comparison | Post-hoc comparison<sup>1</sup> |
| --- | --- | --- | --- | --- | --- | --- | --- | --- |
|  | CUD (n = 44) |  | TUD (n = 18) |  | OUD (n = 16) |  |  |  |
| Demographics |  |  |  |  |  |  |  |  |
| Sex (female / male), *N* (%) | 19 / 25 | (43 / 57) | 8 / 10 | (44 / 56) | 4 / 12 | (25 / 75) | $\chi^2(2) = 1.84, p = .399$ | |
| Site (Berlin / Mannheim), *N* (%) | 18 / 26 | (41 / 59) | 9 / 9 | (50 / 50) | 4 / 12 | (25 / 75) | $\chi^2(2) = 2.27, p = .322$ | |
| Age in years, *M* (± *SD*) | 33.23 | (± 7.95) | 36.39 | (± 9.86) | 40.81 | (± 6.42) | $F(2, 75) = 5.21, p = .008$ | OUD > CUD |
| Educational level, *median* (*IQR*) | 5 | (4–5) | 5 | (3–5) | 4 | (3–5) | $H(2) = 3.37, p = .185$ | |
| Psychometric assessment, *M* (± *SD*) | | | | | | | | |
| Craving (MaCS) | 28.39 | (± 9.11) | 23.50 | (± 7.65) | 27.63 | (± 8.25) | $F(2, 75) = 2.08, p = .132$ | |
| Alcohol use (AUDIT) | 5.20 | (± 4.50) | 3.64 | (± 2.95) | 6.57 | (± 6.65) | $F(2, 75) = 1.32, p = .274$ | |
| Smoking severity (FTND) | 3.21 | (± 1.28) | 4.00 | (± 1.14) | 4.13 | (± 1.45) | $F(2, 75) = 4.03, p = .022$ | OUD > CUD |
| Depression (BDI) | 14.43 | (± 12.15) | 8.56 | (± 7.98) | 10.97 | (± 8.71) | $F(2, 75) = 1.91, p = .156$ | |
| Anxiety State (STAI-S) | 42.79 | (± 12.66) | 40.03 | (± 9.25) | 45.54 | (± 7.44) | $F(2, 75) = 1.05, p = .354$ | |
| Anxiety Trait (STAI-T) | 45.08 | (± 13.74) | 42.48 | (± 9.99) | 43.48 | (± 10.72) | $F(2, 75) = 0.31, p = .733$ | |
| Perceived stress (PSS) | 18.25 | (± 8.91) | 15.55 | (± 5.54) | 18.37 | (± 4.94) | $F(2, 75) = 0.90, p = .410$ | |
| fMRI quality, *M* (± *SD*) | | | | | | | | |
| Head motion (mean FD) | 0.17 | (± 0.06) | 0.19 | (± 0.07) | 0.21 | (± 0.08) | $F(2, 75) = 2.17, p = .122$ | |

<sup>1</sup>Post-hoc pairwise comparisons were Bonferroni-corrected, accounting for the number of group comparisons. Abbreviations: SUD = Substance Use Disorder, CUD = Cannabis Use Disorder, TUD = Tobacco Use Disorder, OUD = Opioid Use Disorder, IQR = interquartile range, MaCS = Mannheim Craving Scale, AUDIT = Alcohol Use Disorder Identification Test, FTND = Fagerström Test for Nicotine Dependence, BDI = Beck Depression Inventory, STAI = State Trait Anxiety Inventory, PSS = Perceived Stress Scale, FD = framewise displacement.

### Connectome-based Predictive Modeling results

Both the positive and negative networks significantly predicted self-reported craving across SUDs (positive network: *rho* = 0.27, *r* = 0.27, *q²* = 0.03, *MSE* = 73.33, *p* = .017; negative network: *rho* = 0.22, *r* = 0.24, *q²* = 0.04, *MSE* = 72.52, *p* = .046). The combined network demonstrated the strongest predictive performance (*rho* = 0.42, *r* = 0.42, *q²* = 0.15, *MSE* = 64.22, *p* = .003; Figure 1A).

**Figure 1.**
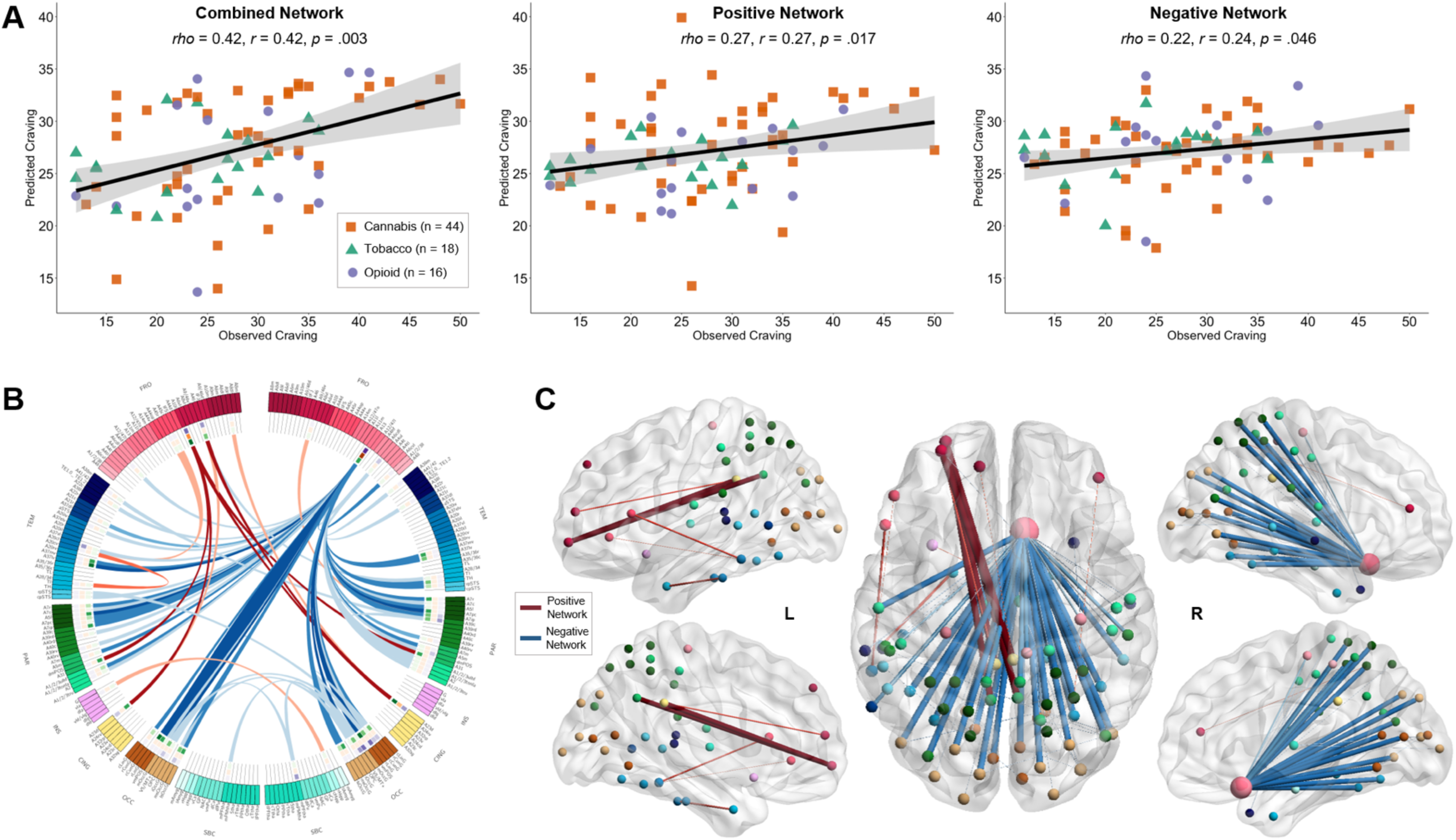
Connectome-based predictive modelling of craving. A) Correlations between true and predicted craving scores using combined, positive, and negative networks across SUD groups (10,000 permutations). B) Connectogram of positive (red) and negative (blue) networks based on Brainnetome Atlas parcellation. Color shadings represent different anatomical subregions based on the Brainnetome Atlas. C) Surface map of craving networks, with larger spheres indicating nodes with higher number of connections.

### Control analyses

Using both a more liberal (*p* < .01) and a more conservative (*p* < .001) threshold for feature selection yielded significant predictions of craving, though performance was lower with the more liberal threshold (Supplementary Table 4).

In addition, accounting for covariates such as substance class, age, sex, site, education, smoking severity, head motion, as well as including all covariates simultaneously, did not alter prediction performance, with all models significantly predicting craving, and none of the models achieving significantly lower prediction performance compared to the original model (Supplementary Table 5).

### Network characterization

The combined craving network consists of 79 edges (0.26% of all edges) connecting 77 nodes across the brain. Of these edges, 12 belonged to the positive network, while 67 were part of the negative network (Figure 1B). The top 5% of nodes with the highest degree were located in the left lateral middle frontal gyrus (BA 10), right rostral lingual gyrus, right middle occipital gyrus, and right medial orbitofrontal cortex (OFC; BA 13). Notably, the right medial OFC emerged as a hub region, accounting for 67.09% of all connections within the identified network (Figure 1C).

Regarding the contribution of large-scale canonical networks, the positive network was composed predominantly of connections within the default mode network (DMN) and the frontoparietal network (FPN) (Figure 2, top row), whereas the negative network mainly comprised connections involving the limbic, somato-motor, visual, and dorsal attention network (DAN) (Figure 2, bottom row).

**Figure 2.**
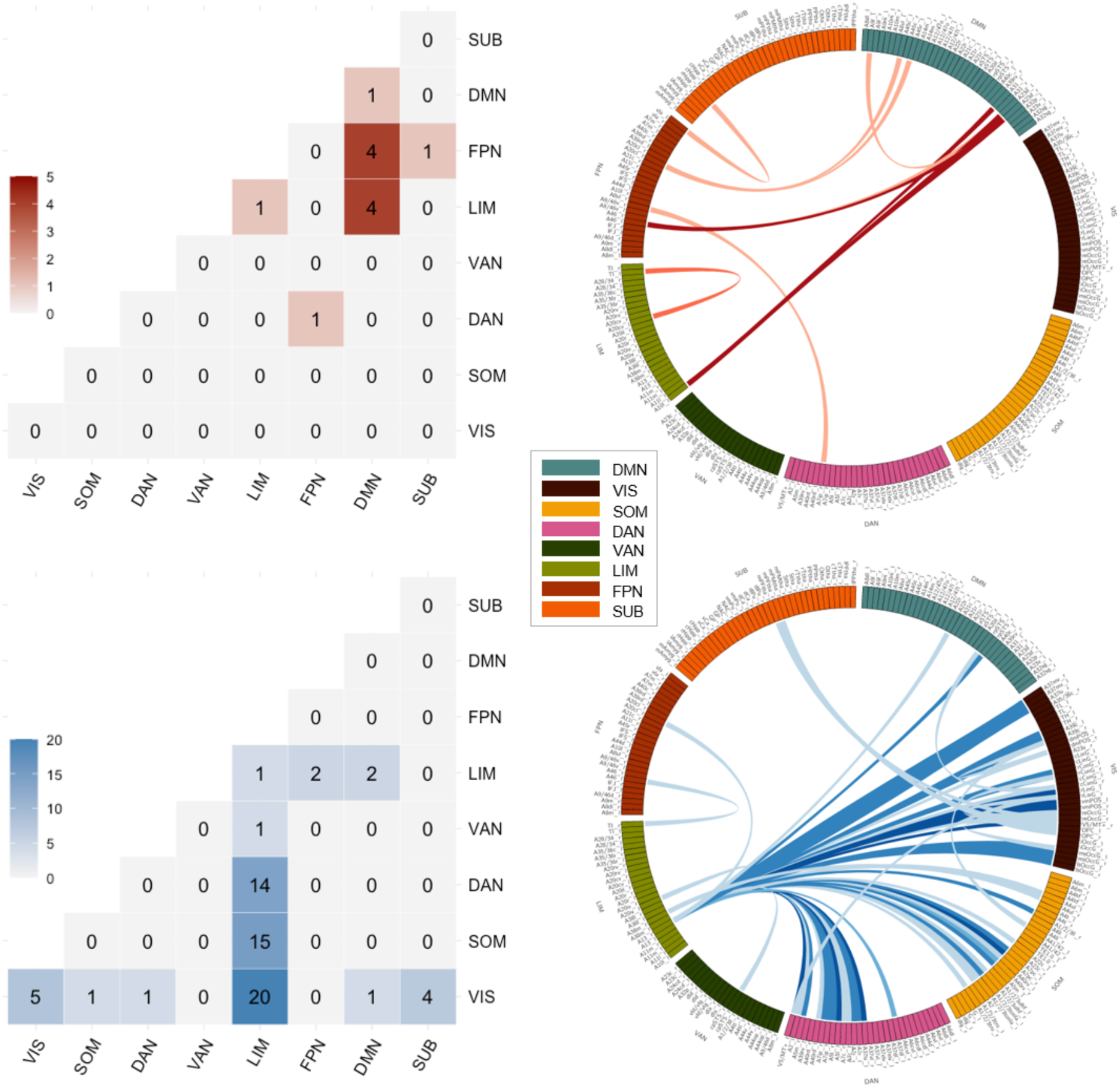
Network-level contribution of the positive network (top row) and negative network (bottom row) for craving prediction. Heatmaps indicate the number of connections between canonical brain networks. Abbreviations: DMN = default mode network, VIS = visual network, SOM = somatomotor network, DAN = dorsal attention network, VAN = ventral attention network, LIM = limbic network, FPN = frontoparietal network, SUB = subcortical network.

### Computational lesion analysis

Node-wise lesioning indicated that the largest drop in prediction performance compared to the full model (*Δrho*) was observed when removing the right medial OFC (BA 13; *Δrho* = 0.41, *z* = 3.26, *p* < .001). Moreover, lesions in the left lateral middle frontal gyrus (BA 10; *Δrho* = 0.21, *z* = 1.78, *p* = .038), the right dorsal posterior cingulate cortex (BA 23; *Δrho* = 0.10, *z* = 2.61, *p* = .004), the left lateral superior parietal lobule (BA 5; *Δrho* = 0.0040, *z* = 1.91, *p* = .028), the bilateral ventromedial parietooccipital sulcus (left: *Δrho* = 0.0017, *z* = 2.17, *p* = .015; right: *Δrho* = 0.0016, *z* = 2.19, *p* = .014), and the left rostral lingual gyrus (*Δrho* = 0.0013, *z* = 1.82, *p* = .034) resulted in significantly lower prediction performance compared to the full model (Figure 3).

**Figure 3.**
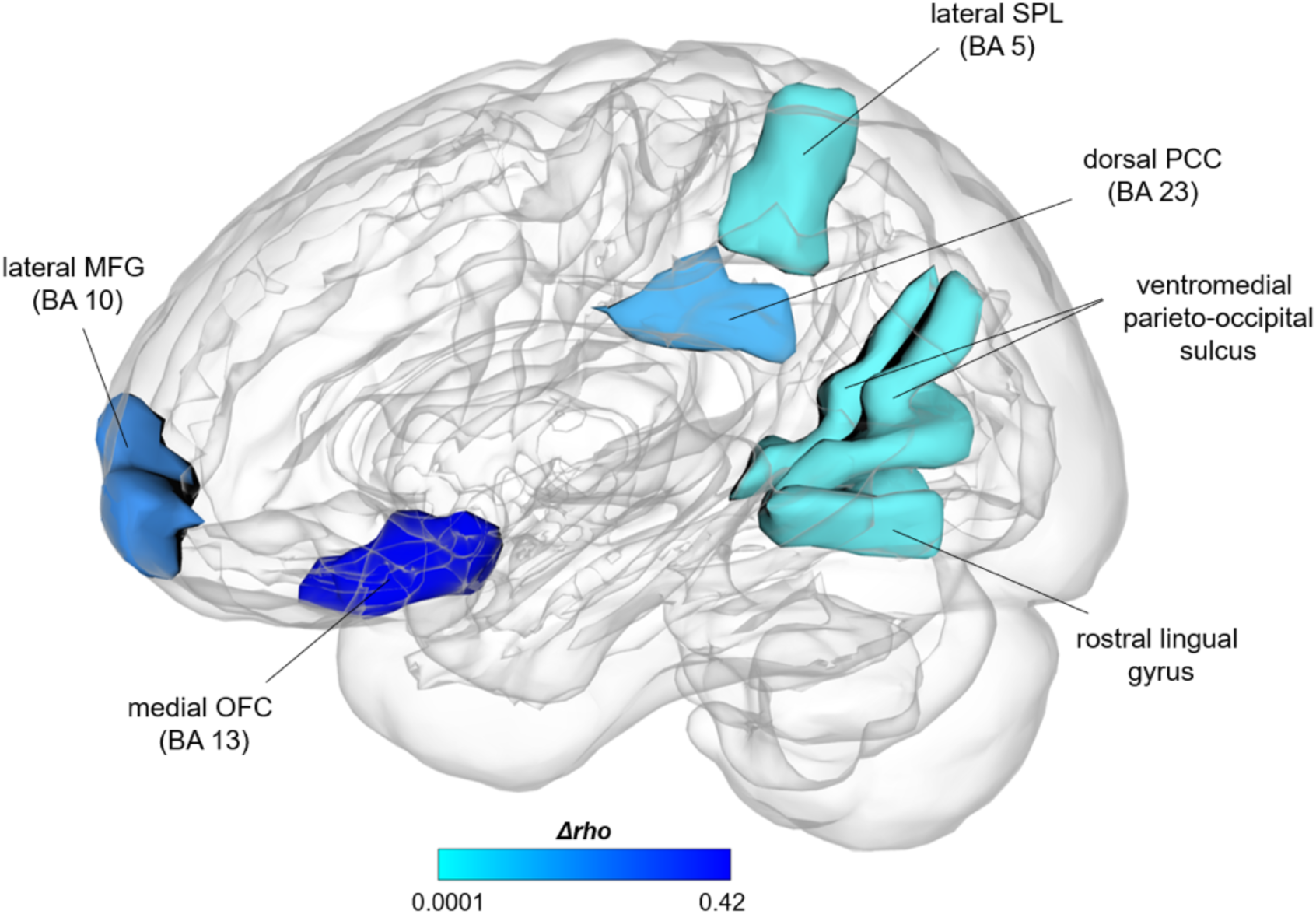
Node-wise virtual lesion analysis showing brain regions whose removal from the network led to a significant reduction in prediction performance compared to the full model (*Δrho*), indicating their importance for craving prediction. Abbreviations: BA = Brodmann area, MFG = middle frontal gyrus, OFC = orbitofrontal cortex, PCC = posterior cingulate cortex, SPL = superior parietal lobule.

Network-wise lesioning demonstrated that excluding the limbic network as well as the DMN led to a significant decline in prediction performance as compared to the full model (Supplementary Table 6).

### External validation I: Alcohol dependence

In an independent sample of patients with alcohol dependence, the identified craving network significantly predicted craving levels on the *Thoughts* subscale of the OCDS-G, reflecting cognitive preoccupation with alcohol (*rho* = 0.33, *r* = 0.28, *p* = .018). Interestingly, the positive network was also significantly associated with this subscale (*rho* = 0.33, *r* = 0.28, *p* = .018), whereas the negative network was not (*rho* = -0.03, *r* = 0.01, *p* = .586) (Figure 4A, top row). In contrast, craving levels on the *Impulse to Act* subscale, reflecting the urge to drink, were significantly predicted by both the overall craving network (*rho* = 0.29, *r* = 0.32, *p* = .033) and particularly by the negative network (*rho* = 0.30, *r* = 0.29, *p* = .027), while the positive network showed no significant association (*rho* = -0.01, *r* = 0.13, *p* = .503) (Figure 4A, bottom row).

**Figure 4.**
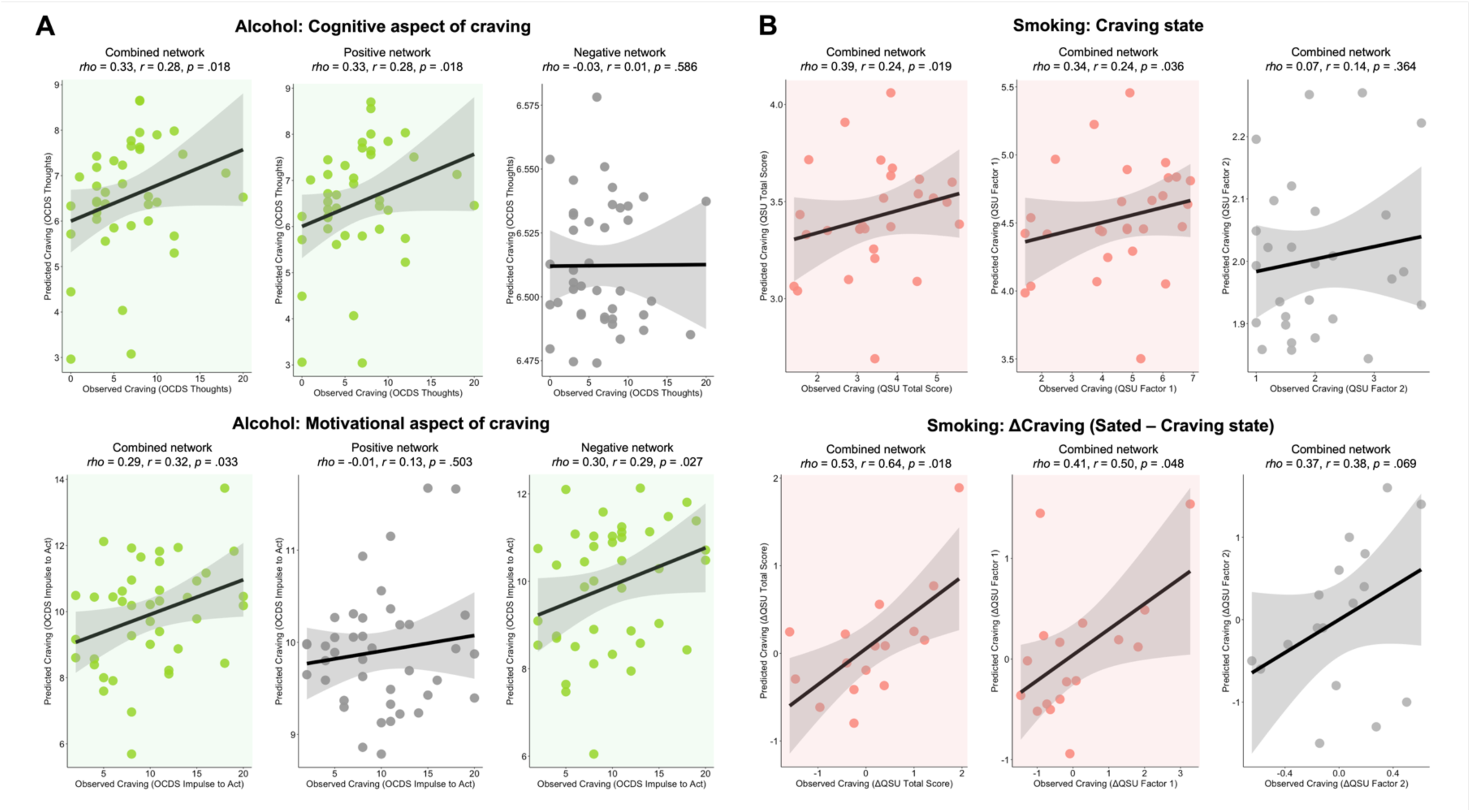
A) External validation I. Top row: In an independent sample of alcohol-dependent patients, the combined network and the positive network predicted cognitive preoccupation with alcohol, indicated by the OCDS-G *Thoughts* subscale. Bottom row: The combined network and the negative network predicted the urge to drink, indicated by the OCDS-G *Impulse to Act* subscale. Shading indicates significant (light green) and non-significant (grey) results. B) External validation II. Top row: In an independent sample of tobacco users, the identified craving network predicted both overall nicotine craving (QSU Total score) and the desire and intention to smoke (QSU Factor 1) after participants had undergone at least 10 hours of smoking abstinence. Bottom row: Changes in functional connectivity strength within the craving network from the sated to the craving state were associated with corresponding changes in overall nicotine craving (QSU Total score) and in the desire and intention to smoke (QSU Factor 1). No significant associations were found for the anticipation of relief from negative affect (QSU Factor 2). Shading indicates significant (light red) and non-significant (grey) results.

### External validation II: Tobacco Use Disorder

Moreover, our identified craving network significantly predicted overall nicotine craving (QSU-G total score: *rho* = 0.39, *r* = 0.24, *p* = .019), as well as the desire and intention to smoke (QSU-G *Factor 1*: *rho* = 0.34, *r* = 0.24, *p* = .036) in an independent sample of tobacco users who were experimentally induced into a craving state (Figure 4B, top row). Importantly, the change in functional connectivity strength within the identified craving network from the sated state to the craving state predicted the corresponding change in overall nicotine craving (QSU-G total score: *rho* = 0.53, *r* = 0.64, *p* = .018) as well as the change in the desire and intention to smoke (QSU-G *Factor 1*: *rho* = 0.41, *r* = 0.50, *p* = .048) (Figure 4B, bottom row). No significant predictions were observed for the anticipation of relief from negative affect (QSU-G *Factor 2*).

## DISCUSSION

The present study used connectome-based predictive modeling (CPM) of resting-state functional connectivity data to predict subjective craving across SUDs. In a transdiagnostic sample of individuals with cannabis, opioid, or tobacco use disorder, we identified a complex craving-related network, with the right medial orbitofrontal cortex (OFC), right dorsal posterior cingulate cortex (PCC), and left lateral medial frontal gyrus emerging as key regions. External validation in alcohol-dependent patients showed that the positive network predicted cognitive aspects of craving, while the negative networks predicted motivational aspects. In smokers, the overall network further predicted both craving intensity after abstinence and changes from a sated to a craving state. Given the lack of differences in clinical comorbidities across SUDs and the robustness of the CPM results to covariate control, the identified network likely reflects a transdiagnostic craving signature rather than substance-specific or general clinical effects. Together, these findings point to a robust functional brain network predictive of subjective craving across SUDs.

Consistent with previous findings [13–16,28], the identified craving network was complex, encompassing multiple regions across different large-scale brain systems. Specifically, it comprised a positive network, where stronger connectivity predicted higher craving, and a negative network, where weaker connectivity predicted higher craving. The positive network was primarily characterized by connections linking the default mode network (DMN) and the frontoparietal network (FPN). The DMN is implicated in internally directed processes such as self-referential thinking, memory integration, and mental simulation of past and future events [29]. Within this network, the right dorsal PCC emerged as a particularly influential region for craving prediction. The FPN, by contrast, is typically associated with cognitive control and executive functions, including goal-directed behavior, working memory, and flexible attentional allocation [30,31]. Consistent with this role, the lateral middle frontal gyrus, a core hub of the FPN [30], was also among the most predictive nodes of craving. In line with our findings, prior CPM studies have likewise implicated links and interactions from and within the FPN [13,15,16,28] and DMN [13–16,28] in the prediction of craving across SUDs and behavioral addictions. Together, our findings suggest that heightened craving may be linked to increased coupling between self-referential and control circuits. Potentially, stronger DMN-FPN connectivity reflects the capture of cognitive control resources, whereby the FPN becomes increasingly engaged with craving-related self-referential states, directing attention toward internal urges and substance-related thoughts [32]. Notably, in our validation sample of patients with alcohol dependence, the positive network was particularly associated with the cognitive preoccupation with alcohol, but unrelated to the motivational urge to drink, further supporting its role in the cognitive dimension of craving.

The negative network, in which weaker connectivity was linked to higher craving, was dominated by the right medial OFC, accounting for 80% of negative network connections. As a core region of the reward circuit [20], the medial OFC supports the appraisal of subjective value that guides decision-making [33]. Dysregulation of OFC circuitry has been consistently linked to SUDs [19] and, more specifically, to craving [13,34,35]. In the present study, higher craving was associated with reduced connectivity between the right medial OFC and regions of the visual, somato-motor, and dorsal attention networks. This pattern suggests that the valuation system becomes less integrated with systems that guide attention and process external cues [33]. As a result, valuation processes may be insufficiently modulated by top-down control or competing sensory input, and instead may become dominated by internally generated reward expectations, thereby contributing to the persistence of craving.

Validation of the identified craving network in patients with alcohol dependence revealed two important insights. First, it demonstrated the generalizability of the network beyond the substances included in the original sample, thereby supporting transdiagnostic views of addiction that emphasize shared circuit-level dysfunctions across different substances [18–20]. Second, validation analyses in patients with alcohol dependence showed that the positive network was more closely associated with the cognitive preoccupation of alcohol, while the negative network captured primarily the motivational urge to drink. Craving is widely recognized as a complex, multidimensional construct encompassing cognitive, emotional, motivational, sensory, and physiological components [36,37]. Our findings suggest that these distinct dimensions of craving may be supported by partially separable neural mechanisms.

In addition, validation in a second external dataset of tobacco users revealed that the combined craving network predicted both absolute craving intensity after experimentally induced abstinence, as well as the change in craving relative to a sated baseline. This is consistent with previous work showing that functional changes within fronto-striatal circuits, including the right orbitofrontal cortex (OFC), were associated with changes in craving in smokers following 12 hours of smoking abstinence [35]. This state sensitivity underscores the potential translational value of the identified craving network. Biomarkers that track intra-individual fluctuations might be particularly promising for clinical applications, such as just-in-time interventions [38].

Our study has several limitations that should be considered when interpreting the results. First, the sample size was modest, highlighting the need for larger and more diverse transdiagnostic samples. Second, craving was assessed via subjective self-reports. Although such instruments are widely used [39] due to their validity [40] and ease of administration, future studies could incorporate more objective measures, such as choice paradigms [41], or complement self-reports with ecological momentary assessments to increase ecological validity [42]. Third, craving is a multifaceted construct [36,37] and was assessed differently across datasets. While such heterogeneity could contribute to discrepancies with prior work, the network’s replication across datasets suggests that the identified pattern may not be strongly tied to a specific craving measure or assessment context. Lastly, while our focus was on transdiagnostic modeling of craving within SUDs, craving has shown to be a central phenomenon also in behavioral addictions [43], eating disorders [44], and compulsive sexual behaviors [40]. While prior research suggests converging neural correlates across these domains [14,45], the craving network identified in our study does not allow conclusions about the neural mechanisms of craving in other addictive behaviors.

In sum, by modeling craving across multiple SUDs, this study identified a transdiagnostic functional brain network predictive of subjective craving. Using a connectome-based machine learning approach, it advances beyond descriptive associations toward individualized prediction, a key step for precision psychiatry [6,7]. Given its centrality in the identified craving network, the right medial OFC might serve as a potential target for intervention, such as neuromodulation approaches [46,47]. Validation in independent samples of alcohol-dependent patients and smokers confirmed the network’s generalizability, its sensitivity to distinct cognitive and motivational dimensions of craving, and its ability to capture intra-individual fluctuations of craving. Future work should assess its predictive utility for longitudinal outcomes such as relapse risk and treatment response.

## DATA AVAILABILITY STATEMENT

The raw data are not publicly available due to privacy and ethical restrictions. Derived data may be provided upon reasonable request to the corresponding author.

## AUTHOR CONTRIBUTIONS

HW, WHS, GE, and RS were involved in the planning and conceptualization of the study. JB, LFE, KE, RON, LW, PR, and NZ performed data acquisition and/or data quality control for any of the datasets used in the study. JB performed preprocessing and data analysis. JB drafted the manuscript. LFE, KE, RON, LW, PR, NZ, GW, PB, RS, GE, WHS, and HW provided critical revision of the manuscript for important intellectual content. All authors critically reviewed the content of the manuscript and approved the final version for publication.

