## Supplementary Information for "Brain Functional Connectivity Signatures of Craving Across Substance Use Disorders: A Transdiagnostic Approach"

**TABLE OF CONTENTS**

Supplementary Table 1. Diagnostic criteria for substance use disorders across subjects

Supplementary Table 2. Sample description of replication sample 1 (*TransAlc*) and comparison with discovery sample (*SysMedSUDs*)

Supplementary Table 3. Sample description of replication sample 2 (*SmokingCraving*) and comparison with discovery sample (*SysMedSUDs*)

Supplementary Table 4. Model performance with different feature selection thresholds

Supplementary Table 5. Model performance when adjusting for covariates

Supplementary Table 6. Network-vise virtual lesioning results

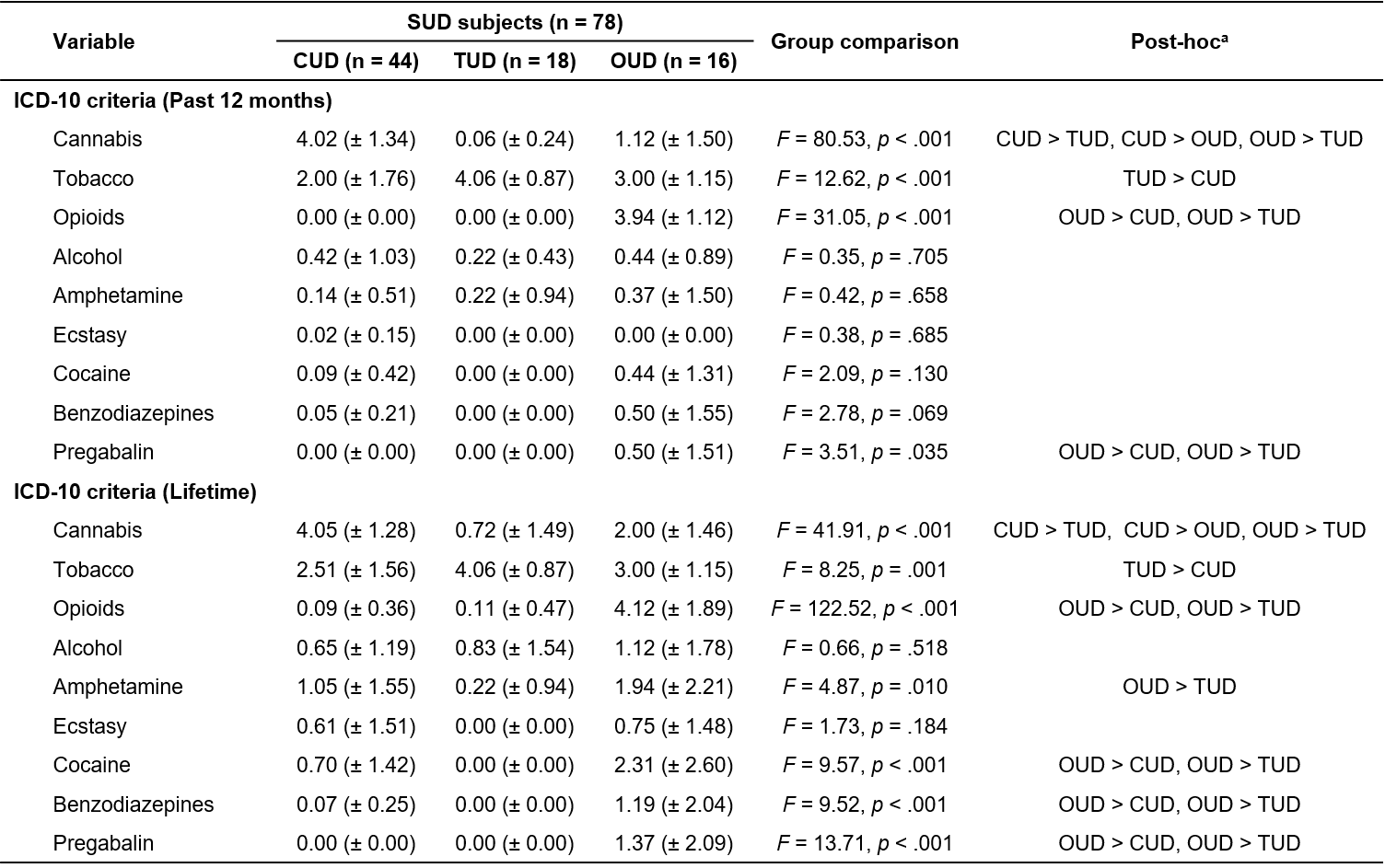
**Supplementary Table 1.** Diagnostic criteria for substance use disorders across subjects

**^a^**Post-hoc pairwise comparisons were Bonferroni-corrected. Abbreviations: SUD = Substance Use Disorder, CUD = Cannabis Use Disorder, TUD = Tobacco Use Disorder, OUD = Opioid Use Disorder.

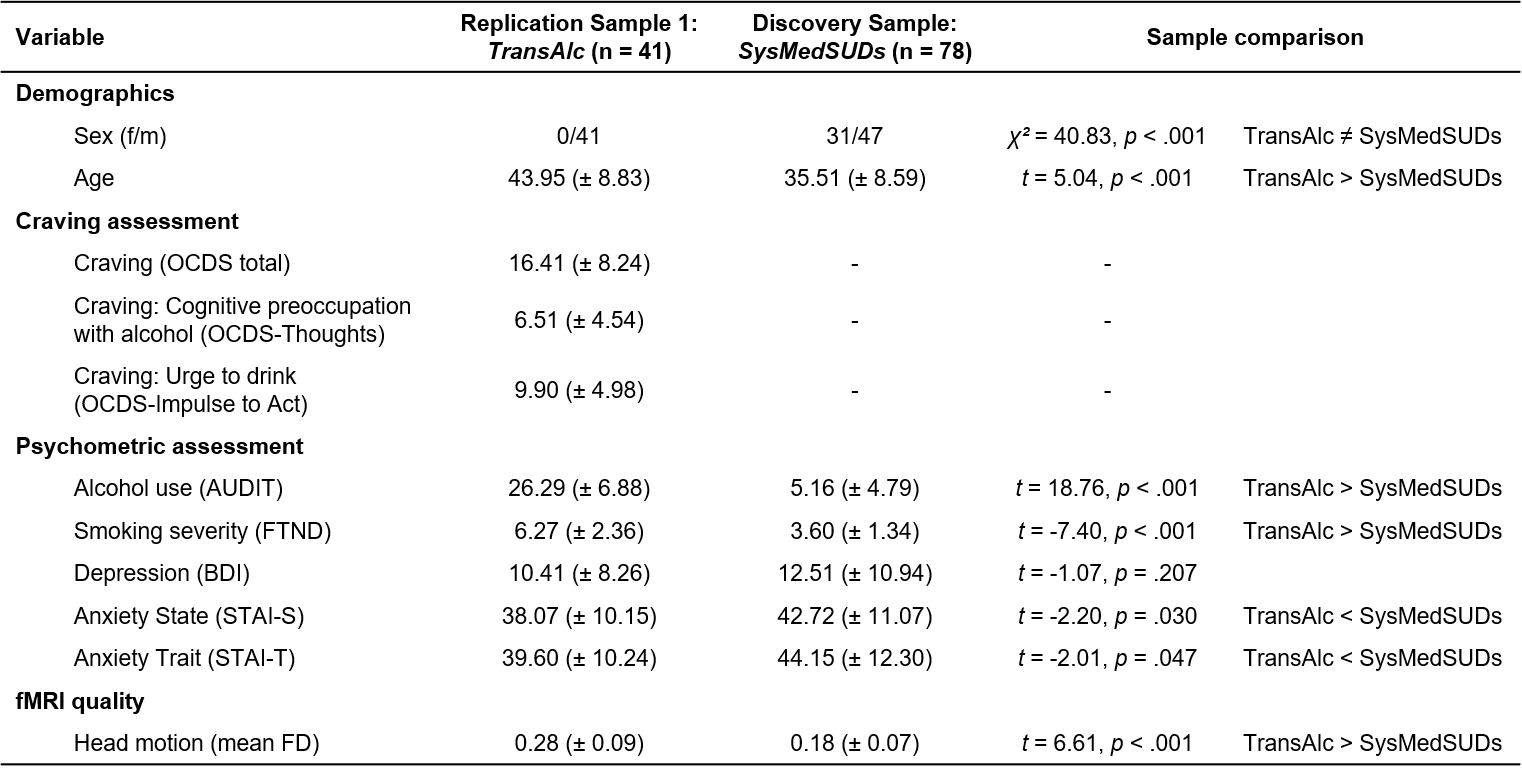
**Supplementary Table 2**. Sample description of replication sample 1 (*TransAlc*) and comparison with discovery sample (*SysMedSUDs*)

Abbreviations: OCDS = Obsessive-Compulsive Drinking Scale, AUDIT = Alcohol Use Disorder Identification Test, FTND = Fagerström Test for Nicotine Dependence, BDI = Beck Depression Inventory, STAI = State Trait Anxiety Inventory, FD = framewise displacement

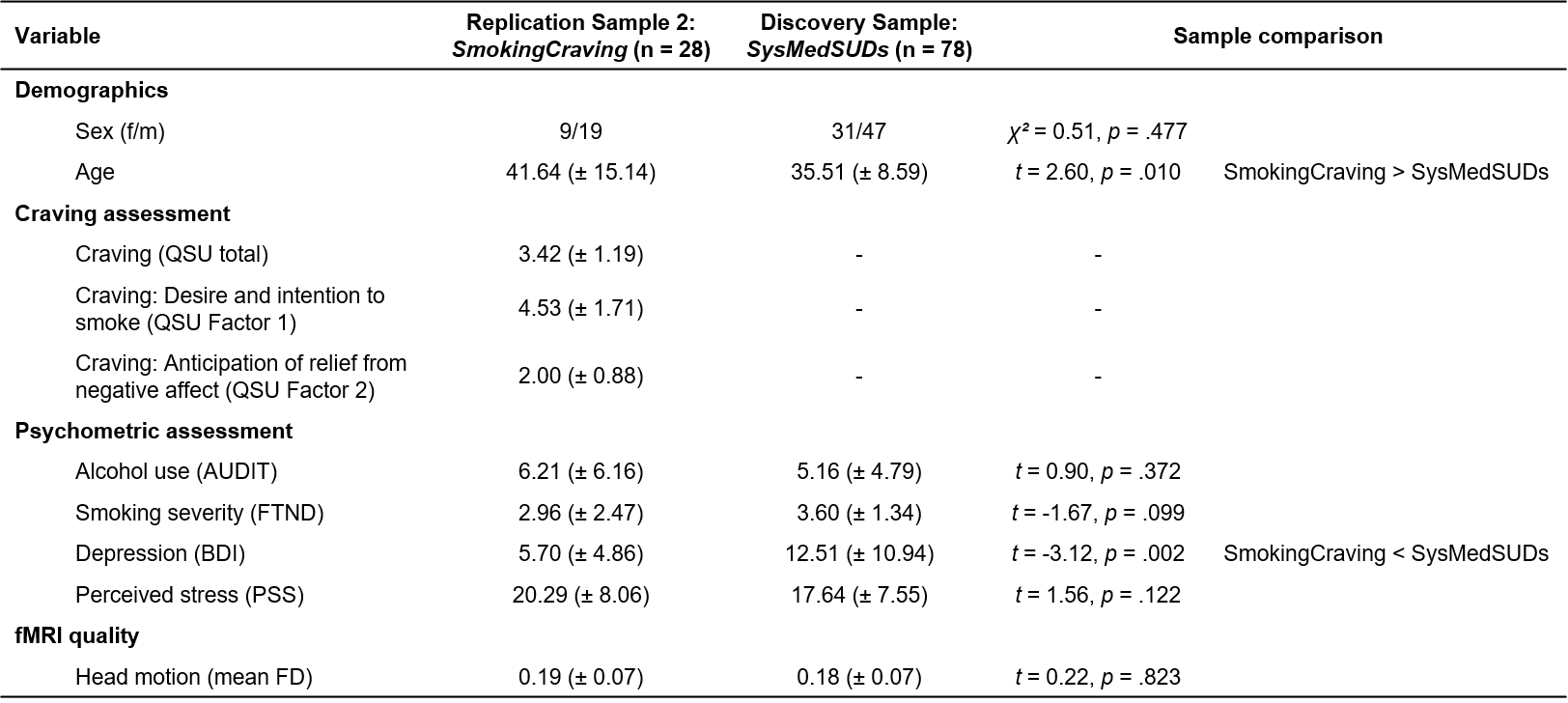
**Supplementary Table 3**. Sample description of replication sample 2 (*SmokingCraving*) and comparison with discovery sample (*SysMedSUDs*)

Abbreviations: QSU = Questionnaire on Smoking Urges, AUDIT = Alcohol Use Disorder Identification Test, FTND = Fagerström Test for Nicotine Dependence, BDI = Beck Depression Inventory, PSS = Perceived Stress Scale, FD = framewise displacement.

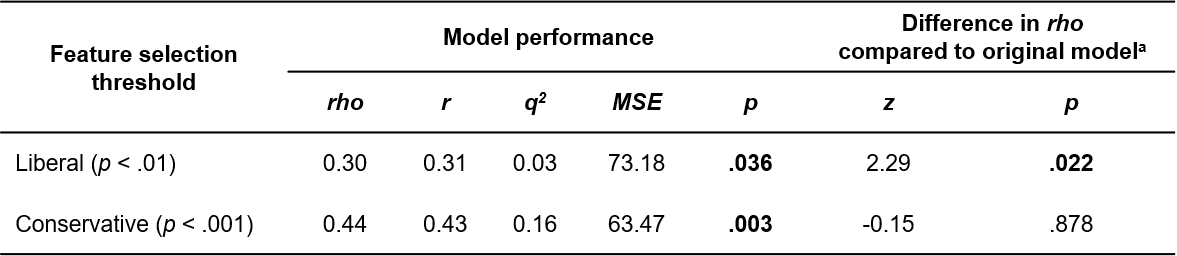
**Supplementary Table 4**. Model performance with different feature selection thresholds

Note: ^a^Difference in prediction performance (rho) between the more liberal or conservative model and the original model reported in the main manuscript based on Steiger‘s z-test.

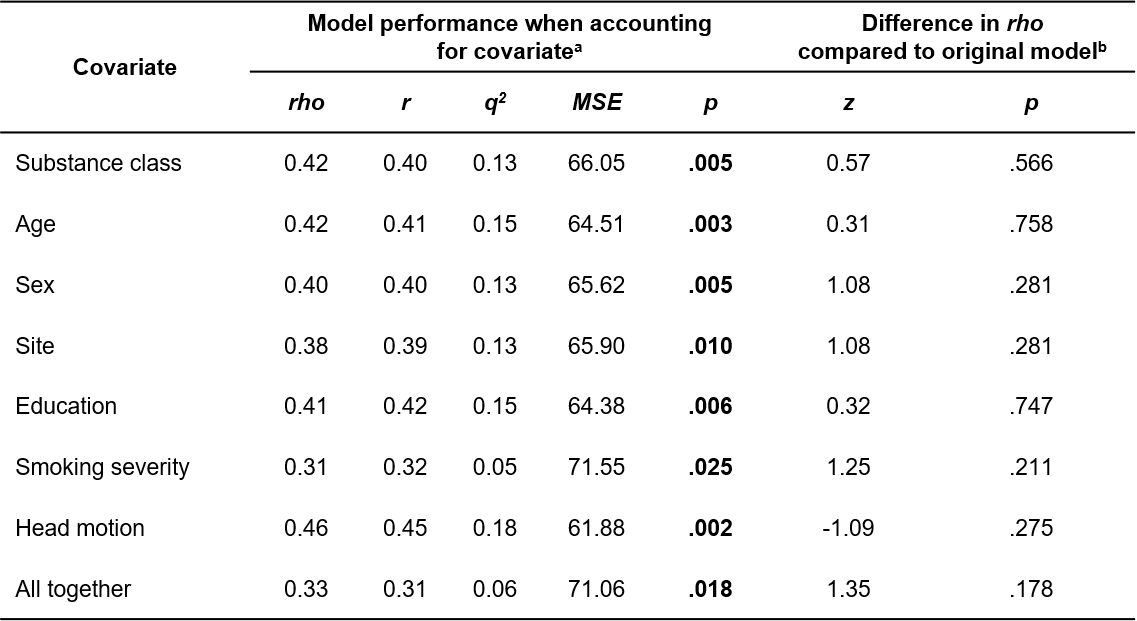
**Supplementary Table 5**. Model performance when adjusting for covariates

Note: ^a^Model performance when accounting for respective covariate using partial correlation during feature selection. Bold values indicate significant prediction of craving. ^b^Difference in prediction performance (*rho*) between the covariate-adjusted model and the original, unadjusted model reported in the main manuscript based on Steiger‘s z-test.

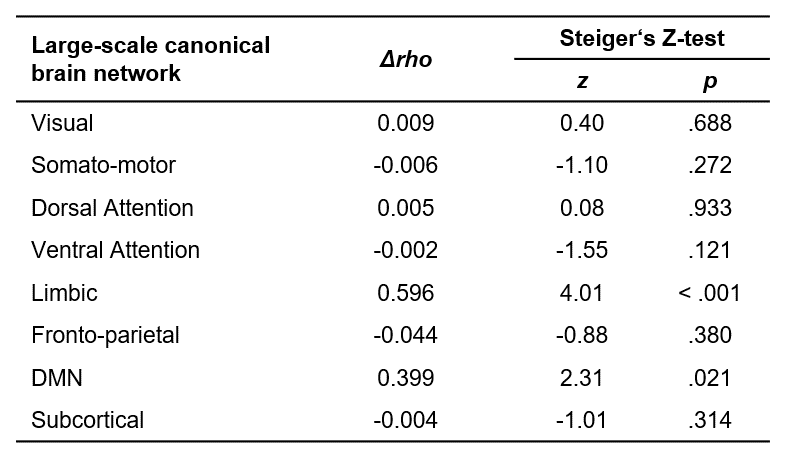
**Supplementary Table 6**. Network-wise virtual lesioning results

Note: *Δrho* indicates the difference in prediction performance between the full network and the lesioned network.
